# Dynamics of blood cells during a routine laboratory examination

**DOI:** 10.1101/2023.01.21.525013

**Authors:** Mesfin Asfaw Taye

## Abstract

Centrifugation is a commonly performed laboratory procedure that helps to separate blood cells such as red blood cells *RBCs*, white bood cells *WBCs*, and platelets from plasma or serum. Although centrifugation is a routine procedure in most medical laboratories, factors that affect the efficacy of the centrifugation process have never been studied analytically. In this paper, we examine the effect of centrifugation time on the efficacy of the centrifugation process by studying the dynamics of blood cells via the well-known Langevin equation or equivalently, by solving the Fokker-Plank equation. Our result depicts that the speed of the centrifuge is one of the determinant factors concerning the efficacy of the centrifugation process. As angular speed increases, centrifugal force increases and as a result, the particles are forced to separate from plasma or serum. The room temperature also considerably affects the dynamics of the sample during centrifugation. Most importantly, the generation of heat during centrifugation increases the temperature within a centrifuge, and as a result, not only the stability of the sample but also the mobility of analyse is affected. We show that as the temperature within the centrifuge intensifies, the velocity of the cells as well as the displacement of the cells in the fluid increases. We then study the dynamics of the whole blood during capillary action where in this case the blood flows upward in a narrow space without the assistance of external forces. Previous investigations show that the height that the fluid rises increases as surface tension steps up. The viscosity of the fluid also affects the capillary action but to date, the dependence of the height on viscosity has never been explored due to the lack of a mathematical correlation between the viscosity of blood and surface tension [1]. In this work, we first examine the correlation between surface tension and viscous friction via data fitting. Our result exhibits that the viscosity of the blood increases linearly as surface tension increases. The mathematical relation between the height and viscous friction is derived. It is shown that the height of the blood that rises in the capillary increases as the viscous friction intensifies. As the temperature of the room steps up, the height also decreases. The dependence of erythrocytes sedimentation rate on surface tension is also studied. The results obtained in this work show that the erythrocyte sedimentation rate ESR increases as surface tension steps down.

**PACS numbers:** Valid PACS appear here

## I. INTRODUCTION

Medical laboratory examinations are vital since these examinations help to diagnose any abnormalities and treat a patient based on observed results. Particularly, blood tests are routinely performed to evaluate any abnormal conditions. Most of these blood works require sedimentation either via centrifugation or gravity [1]. To understand the factors that affect the outcome of routine examinations, several studies have been done [2–9]. Most of these works have focused on exploring the dynamics of whole blood, erythrocytes (RBCs), leukocytes (WBCs), and thrombocytes (platelets) [10].

Particularly, centrifugation is one of the commonly performed laboratory procedure that helps to separate blood cells such as *RBCs, WBCs*, and platelets from plasma or serum [1, 11]. As a common procedure, when blood is centrifuged for a few minutes, the centrifugal force separates the blood cells by leaving the plasma or the serum at the top. In order to understand the factors that affect the efficacy of the centrifugation process, several studies have been done [11–13]. Most recently Minder.et.al [1] studied factors that affect the sedimentation rate of the cells during centrifugation. Most of these experimental studies depict that centrifugation time, temperature, length of the test tube, and speed of the centrifuge are the determinant factors regarding the efficacy of the centrifugation process. Despite several experimental considerations, factors that affect the efficacy of the centrifugation process have never been studied analytically. However analytical solutions are not only important for reconfirming the experimental results but also provide further quantitative powerful predictions.

A hematocrit test is also part of clinical laboratory examinations and measures the percentage of erythrocytes in comparison to whole blood. Using a capillary tube, a blood sample is collected. Via capillary action, blood flows upward in a narrow space without the assistance of external forces. The height that the blood rises is significantly affected by surface tension, viscosity, temperature, and test tube inclination [14]. Previous investigations show that the height that the fluid rises increases as surface tension increases [15]. In the past, for non-Newtonian fluids, Pelofsky [16, 17] has studied the relation between surface tension and viscosity. His empirical equation depicts that surface tension increases as viscosity increases. The effect of sample temperature on the dynamics of whole blood has been explored. The in vivo experiment by Shinozaki *et. al*. [18] indicates that as temperature inclines, blood refill time decreases. The experimental observation by She *et. al*. shows that increasing temperature results in lower capillary pressures (lower capillary height) [19]. The viscosity of blood also affects the capillary action but to date, the dependence of the height on viscosity has never been explored due to the lack of a mathematical correlation between the viscosity of blood and surface tension.

Furthermore, the erythrocyte sedimentation rate (ESR) is one the most common clinical laboratory examination [20]. The erythrocyte sedimentation rate (ESR) often measures how fast a blood sample sediments along a test tube in one hour in a clinical laboratory [10]. This analysis is performed by mixing whole blood with an anticoagulant. The blood is placed in an upright Wintrobe or Westergren tube and allowed to sediments for an hour. The normal values of the ESR vary from 0-3mm/hr for men and 0-7 mm/hr for women [21–23]. High ESR is associated with diseases that cause inflammation. In the case of inflammatory disease, the blood level of fibrinogen becomes too high [23, 24]. The presence of fibrinogen forces the RBCs to stick to each other and as a result, they form aggregates of RBC called rouleaux. As the mass of the rouleaux increases, the weight of the rouleaux dominates the vicious friction and as a result, the RBCs start to precipitate. The temperature of the laboratory (blood sample) also significantly affects the test result [25]. As the temperature of the sample steps up, the ESR increases. In the past, a mathematical model was developed by Sharma .*et. al* to study the effect of blood concentration on the erythrocyte sedimentation rate [26]. Later the effect of the concentration of nutrients on the red blood cell sedimentation rate was investigated in the work [27]. More recently, the sedimentation rate of RBC was explored via a model that uses Caputo fraction derivative [28]. The theoretical work obtained in this work was compared with the sedimentation rate that was analyzed experimentally. More recently, the experimental and analytical studies indicate that red blood cells sediment as a percolating gel and the dynamics of these cells can be studied using equations for porous networks [2–6] All of these experimental and theoretical works exposed the factors that affect the sedimentation rate of RBCs. However, the effect of surface tension as well as the tilt in the test tube on the sedimentation rate has never been explored.

The purpose of this theoretical work is to explore the factors that affect the dynamics of blood during in vivo and in vitro experiments. As discussed before, the factors that affect the efficacy of the centrifugation process have been only studied experimentally. In this work, via analytical solutions, we reconfirm the previous experimental results as well as provide additional quantitative powerful predictions. We then study the dynamics of whole blood during capillary action. We first examine the correlation between surface tension and viscous friction via data fitting. The mathematical relation between height and viscous friction is also derived. The dependence of the erythrocyte sedimentation rate (ESR) on model parameters is explored analytically. Since the effect of surface tension on sedimentation rate has never been considered before, in this paper we study the role of surface tension on the sedimentation rate analytically.

Since blood cells are microscopic, their dynamics can be modeled as a Brownian particle walking in a viscous medium. As blood is a highly viscous medium, the chance for the blood cells to accelerate is negligible and the corresponding dynamics can be studied via Langevin equation or Fokker Planck equation [29–35]. Solving the Fokker-Planck equation analytically (the Langevin equation), the dynamics of blood cells as a function of the model parameters can be explored. Because our study is performed by considering real physiological parameters, the results obtained in this work non only agree with the experimental observations but also help to understand most hematological experiments that are conducted in vitro or in vivo. The alternative approaches depicted in the works [2–6] also indicate that since red blood cells at high fraction volume form a gel, their dynamics can be uniquely studied via equations for porous networks.

The analytical results that are obtained in this theoretical work depict that the speed of the centrifuge is one of the determinant factors concerning the efficacy of the centrifugation process. As the angular speed increases, the centrifugal force steps up and as a result, the particles are forced to separate from the plasma or serum. As shown in Fig. 1 [36], the size of the WBC is considerably large in comparison with the RBC. Since the platelet has the smallest size, its velocity and displacement along the test tube are significantly small. As a result, RBCs separate far more than platelets showing that to avoid the effect of clotting factors, one has to increase the centrifugation time. The height that the blood rises up along a capillary tube increases as the viscous friction steps up. As the temperature of the room steps up, the height also decreases.

**FIG. 1:**
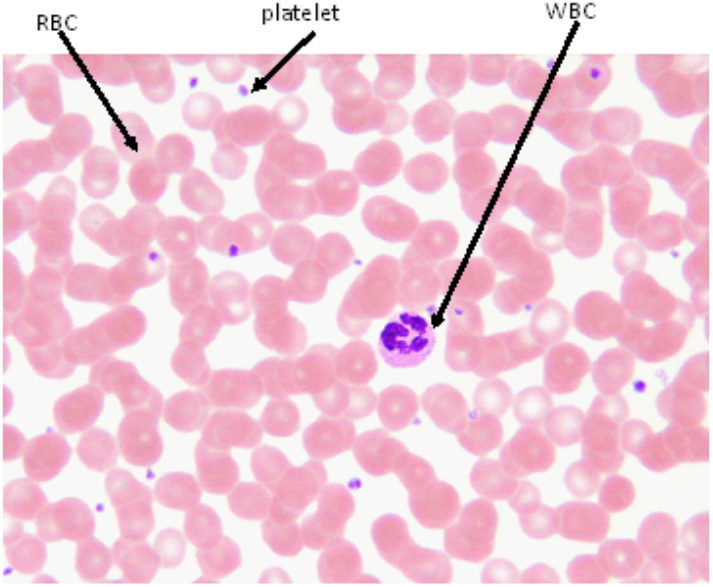
A blood smear that shows the composition of RBCs, WBCs, and platelets [36].

It is important to emphasize that the present study can potentially impact the current research field by providing a new powerful prediction and also by serving as a guide on how to correct false negative or false positive results. As discussed before, the room temperature considerably affects the dynamics of the sample during centrifugation. Most importantly, the generation of heat during centrifugation steps up the temperature within a centrifuge, and as a result, not only the stability of the sample but also the mobility of analyse is affected. The effect of temperature near the test tube causes difficulties in the current experimental procedure by inducing additional false-positive results. In this regard, developing a mathematical model and exploring the model system using the powerful tools of statistical mechanics provides insight as well as guidance regarding the dynamics of RBCs in vitro. Once calculating the exact analytic results, one can correct the extra false positive or negative results. The developed model system also provides a clue about how the dynamics of blood during a routine lab examination are affected by other model parameters.

The rest of the paper is organized as follows. In section II, we present the model system. In section III, we study the factors that affect the efficacy of the centrifugation process. The dynamics of whole blood during capillary action is studied in section IV. In section V, the dependence of erythrocytes sedimentation rate on model parameters is studied. Section VI deals with the summary and conclusion.

## II. THE MODEL

Since RBC is microscopic in size, its dynamics (in vitro) can be modeled as a Brownian particle that undergoes a biased random walk on one-dimensional test tube. In the routine hematology test, the erythrocyte dynamics is also affected by gravitational force

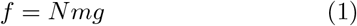

and centrifugal force

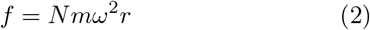

where *g* = 9.8*m/s*^2^ is the gravitational acceleration. *N* denotes the number of blood cells that forms rouleaux. *ω* denotes the angular speed of the centrifuge (see Fig. 2) and *r* designates the radius of the shaft. The speed of clinical centrifuge *ω* varies from 200 rpm (21*rad/s*) and 21000 rpm (2198*rad/s*). The mass of the red blood cells *m* = 27*X*10^−15^ kg. Since platelets are only 20 percent of the size of *RBC*, we infer the mass of the platelets to be *m* = 5.2*X*10^−15^ kg. The average size of RBC and platelets is given as *r*^*′*^ = 4*X*10^−6^ and *r*^*′*^ = 7.5*X*10^−7^ meter, repectively. The normal value of RBC on average varies from 5*X*10^6^ − 6*X*10^6^*/mm*^3^ [37–39].

**FIG. 2:**
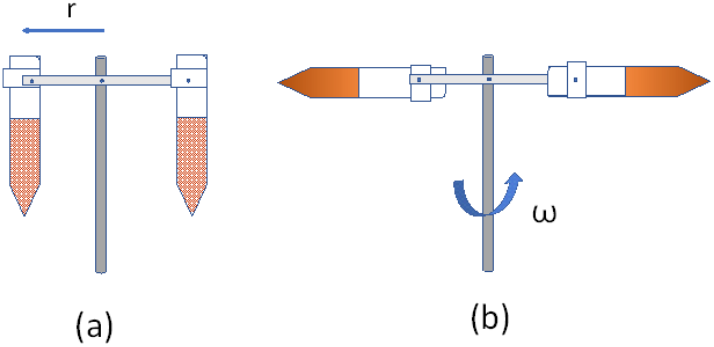
(a) Blood sample before centrifugation. (b) Blood sample during centrifugation. A centrifuge in a clinical laboratory is used to separate blood cells such as *RBCs, WBCs*, and platelets from the plasma or serum. Here *r* designates the radius of the shaft and *ω* denotes the angular speed of the centrifuge.

### Underdamped case

The dynamics of the blood cells in vitro is governed by the well-known Langevin equation

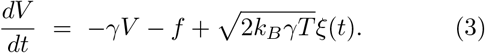

The random noise *ξ*(*t*) is assumed to be Gaussian white noise satisfying the relations ⟨*ξ*(*t*)⟩ = 0 and ⟨*ξ*(*t*)*ξ*(*t*^*′*^)⟩ = *δ*(*t* − *t*^*′*^). The viscous friction *γ* and *T* are assumed to be spatially invariant along with the medium. For a non-Newtonian fluid such blood, it is reasonable to assume that when the temperature of the blood sample increases by 1 degree Celsius, its viscosity steps down by 2 percent [40] as 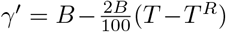 where *B* = 4*X*10^−3^*kg/ms* is the dynamical viscosity of blood at a room temperature (*T*^*R*^ = 20 degree Celsius) and *T* is the temperature [34]. On the other hand, from Stokes’s theorem, the viscosity *γ* = 6*r*^*′*^*πγ*^*′*^. Here *k*_*B*_ = 1.38*X*10^−23^*m*^2^*kgs*^−2^*K*^−1^ is the Boltzmann constant.

At this point, it is important to point out that red blood cell has the shape of a biconcave discoid and undergoes complicated three-dimensional dynamics. As a matter of fact, the shape of the cell dictates the dynamics of the cells both in vivo and vitro. For instance, in the case of sickle cell disease, the sedimentation of the RBC has a lower rate since the abnormal shape of the cell prevents rouleaux formation. Although it is important to consider the biconcave discoid shape of RBC, theoretically (either analytically or numerically) studying the dynamics is quite a cumbersome task. In this work, as an approximation, we modeled the cell as a perfectly spherical object so that one can use Stokes theorem to find the viscous friction.

Alternatively, Eq. (3) can be rewritten as a Fokker-Plank equation

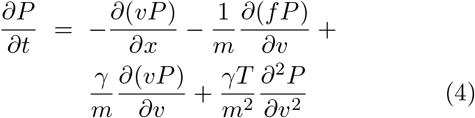

where *P* (*x, v, t*) is the probability of finding the particle at particular position *x*, velocity *v* and time *t*.

For convenience, Eq. (4) can be rearranged as

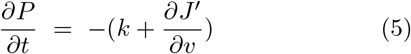

where

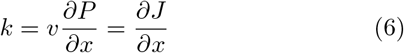

and

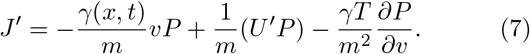

From Eqs. (6) and (7), one gets

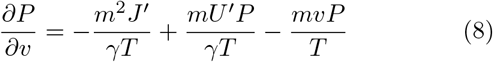

and

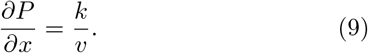

After some algebra, the expression for the probability distribution *P* (*v, t*) is given as

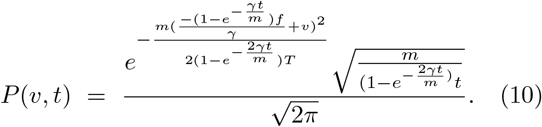

The velocity of the cell can be evaluated as

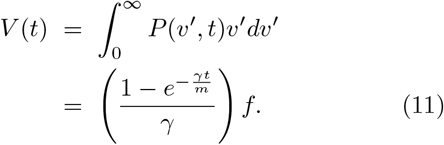

Once the velocity of the cell is calculated, the position of the cell is then given by

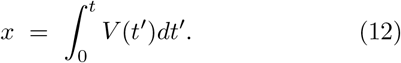

Here one should note that the solutions for overdamped and underdamped cases are not new since such cases are presented by a well-known Ornstein-Uhlenbeck process.

### Overdamped case

Blood is a highly viscous medium, as the result, the chance for the cells to accelerate is negligible. One can then neglect the inertia effect and the corresponding dynamics can be studied via the Langevin equation

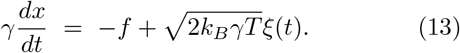

It is worth mentioning that RBC has a disk diameter of 8 microns which is large. However, historically, Brownian motion was first observed through a microscope by looking at pollen (much larger than RBC), and after a few years, Albert Einstein modeled the motion of the pollen and wrote the dynamical equation that describes the motion of this particle. One can also use Langevin equation to exactly derive Einstein’s equation. In other words, Brownian motion occurs at a microscopic and millimeter scale and consequently, the Langevin equation effectively describes the motion of these particles. The Langevin equation better works when the size of the particle is too small. However, the Langevin equation still remains valid to describe the dynamics of the RBC as long as its size is much larger than the dimension of the particles of fluid that the RBC placed.

One can also write Eq. (13) as a Fokker-Plank equation

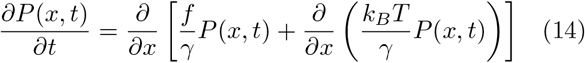

where *P* (*x, t*) is the probability density of finding the particle (the cell) at position *x* and time *t*.

To calculate the desired thermodynamic quantity, let us first find the probability distribution. After imposing a periodic boundary condition *P* (0, *t*) = *P* (*L, t*), we solve Eq. (14). After some algebra, one finds the probability distribution as

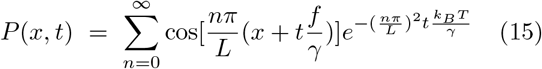

where *T* is the temperature of the medium. For detailed mathematical analysis, please refer to my previous work [34]. Next we use Eq. (15) to find the current, velocity as well as position of the particle. The particle current is then given by

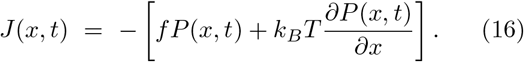

After substituting *P* (*x, t*) shown in Eq. (15), one can find the velocity of the cells at any time as

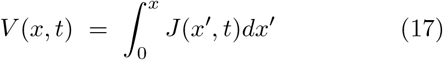

while the position of the cells can be found via

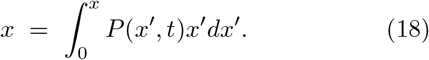

The fact that blood is a highly viscous medium (since *γ* is considerably high), the numerical value of velocity calculated via Eq. (11) is approximately the same as the velocity calculated via Eq. (17). At steady state (in long time limit), the velocity (Eqs. (11) and (17)) approach *V* = *f/γ*. One should also note that the diffusion constant for the model system is given by 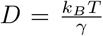. This equation is valid when viscous friction is temperature-dependent showing that the effect of temperature on the mobility of the cells is significant. When temperature increases, the viscous friction gets attenuated and as a result the diffusibility of the particle increases. Various experimental studies also showed that the viscosity of the medium tends to decrease as the temperature of the medium increases [41]. This is because increasing the temperature steps up the speed of the molecules, and this in turn creates a reduction in the interaction time between neighboring molecules. As a result, the intermolecular force between the molecules decreases, and hence the magnitude of the viscous friction decreases.

## III. THE DYNAMICS OF BLOOD CELLS DURING CENTRIFUGATION

As a common standard practice, in a clinical laboratory, centrifugation is vital to separate the blood cells such as *RBCs, WBCs*, and platelets from the plasma or serum. When blood is first mixed with an anticoagulant and allowed to be centrifuged for a few minutes, the blood cells separate from the plasma. In the absence of anticoagulant, the blood clots, and when it is centrifuged, the blood cells segregate from the serum. In this section, we study the factors that affect the efficacy of the centrifugation process analytically. We show that the centrifugation time, temperature, the length of the test tube, and the speed of the centrifuge are the determinant factors with regards to the efficacy of the centrifugation process.

First, let us examine the effect of the centrifugation time on the efficacy of the centrifugation process. This can be investigated by tracking the dynamics of the blood cells during centrifugation. The dynamics of the cells in vitro is governed by the well-known Langevin equations (3) or (13). Equivalently, by solving Fokker-Plank equations (4) or (14), the information regarding the mobility of the RBCs can be extracted. In a medical laboratory, the standard test tube has a length of *L* = 150 *mm* and the cells separate from the plasma or serum at the result of the centrifugal force *f* = *Nmω*^2^*r* where *N* denotes the number of cells that form a cluster while *m* designates the mass of *RBCs* or platelets. Since *WBCs* are heavy, they move way before *RBCs*, and as an approximation one can disregard their dynamics.

Exploiting Eqs. (11) or (17) one can see that the velocity of the RBC steps up and saturates to a constant value (see Fig. 3a). In the figure, we fix the parameters as *N* = 1, *T* = 22 degree celsius, *r* = 0.1*m* and *L* = 0.1*m*. Via Eqs. (12) or (18), one can track the position of RBC along the test tube during the centrifugation processes. As depicted in Fig. 3b, the particle walks away towards the bottom of the test tube as time steps up. Unlike serum, plasma contains platelets and this indicates that more configuration time is needed for plasma since platelets are small in size.

**FIG. 3:**
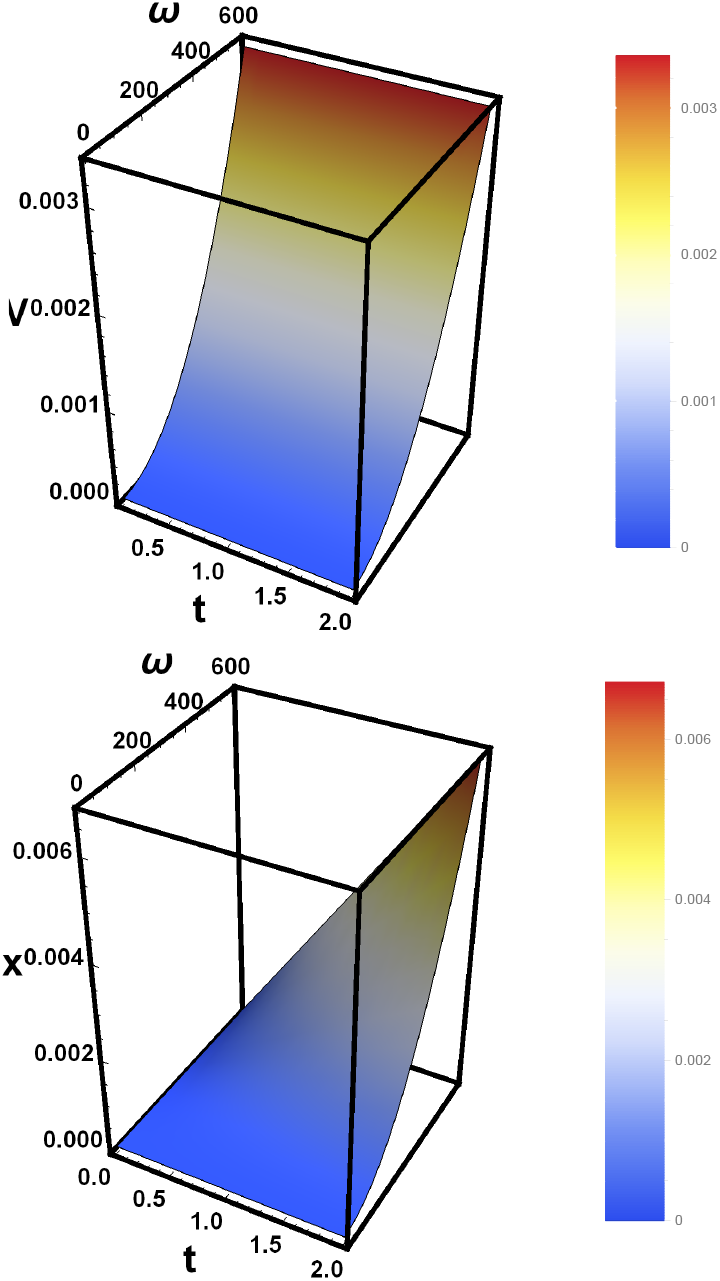
(a) The velocity (*V* (*m/s*)) of RBC as a function of time *t* (in seconds) and *ω* for a single RBC *N* = 1, *T* = 22 degree celsius, *r* = 0.1*m* and *L* = 0.1*m*. (b) The sedimentation displacement (*x*(*m*)) of RBC as a function of time *t* (in seconds) and *ω* for fixed *N* = 1, *T* = 22 degree celsius, *r* = 0.1*m* and *L* = 0.15*m*.

The speed of the centrifuge also affects the efficacy of the centrifugation process. As the angular speed increases (see Eq. (2)), the centrifugal force steps up and as result, the particles are forced to separate from the plasma or serum. Depending on the size and fragility of the analyse, the centrifugation speed should be adjusted. To increase the efficacy of the centrifugation process, as the size of the particle decreases, the centrifugation speed should step up. The dependence of the speed of the particle on angular speed is explored. As depicted in Fig. 3a, the velocity of the RBC steps up monotonously as the angular speed steps up. One can also track the position of RBC along the test tube as a function of angular speed. As shown in Fig. 3b, the particle moves towards the bottom of the test tube as angular speed steps up.

The length of the test tube affects the effectiveness of the centrifugation process. As the length of the test tube steps, the centrifugal force increases. The room temperature considerably affects the dynamics of analyse during centrifugation. Most importantly, the generation of heat during centrifugation increases the temperature within a centrifuge and as a result, not only the stability of the sample but also mobility of analyse is affected. The effect of temperature near the test tube is unavoidable and causes difficulties in the current experimental procedure by inducing additional false-positive results. In this regard, developing a mathematical model and exploring the model system using the powerful tools of statistical mechanics provides insight as well as guidance regarding the dynamics of RBCs in vitro.

The role of temperature on the mobility of the cells during centrifugation can be appreciated by analyzing Eq. (11). Substituting Eq. (2) into Eq. (11), one gets

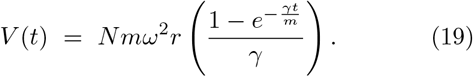

The viscosity *γ* of the blood is the function temperature as 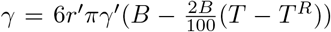 where *r*^*′*^ is the radius of the cells. This implies the velocity *V* (*t*) and position *x* are also the function of temperature. From Eq. (19) one can see that as the centrifuge temperature increases, the velocity of the cells as well as the displacement of the cell in the fluid increases as shown in Figs. 4a and 4b. As discussed before, the effect of temperature can be corrected by increasing the centrifugation time. However, a longer centrifugation time might lead to a considerable amount of heat generation. This, in turn, may cause the red blood cells to lyse. Prolonged centrifugation at high speed also causes structural damage to the cells and as a result, hemolysis occurs.

**FIG. 4:**
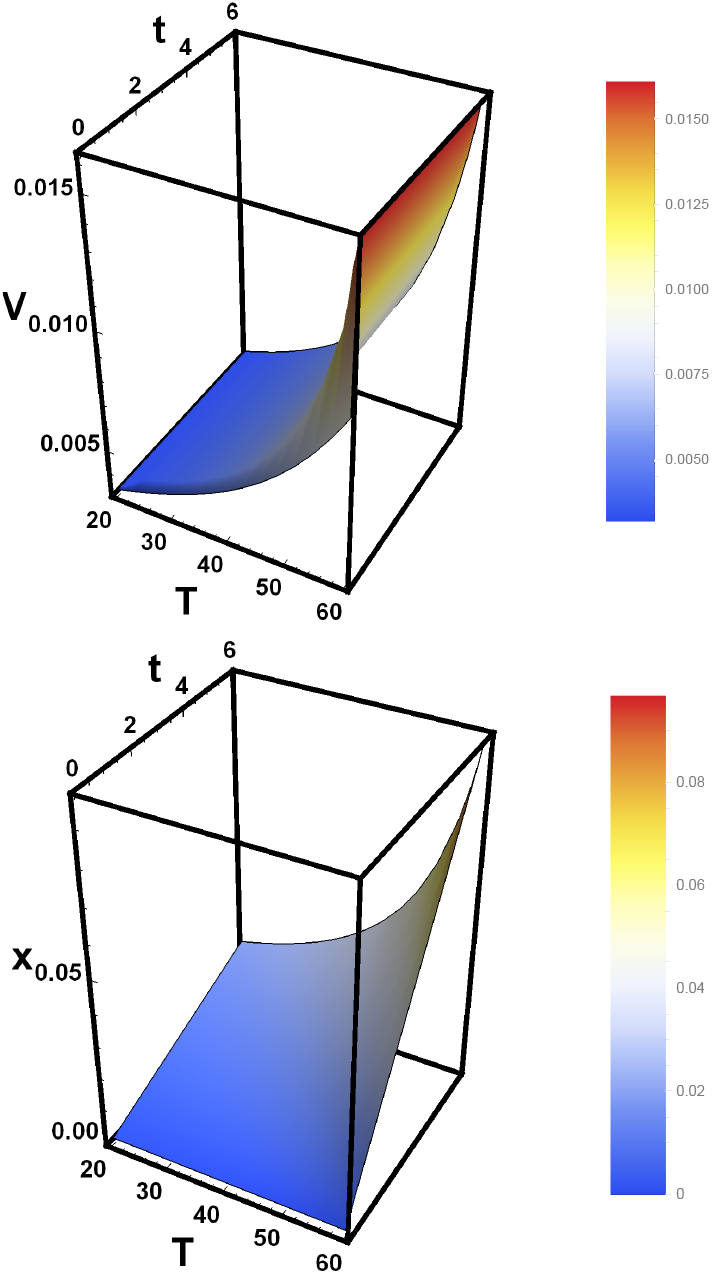
(a) The velocity (*V* (*m/s*)) of RBC as a function of time *t* (in seconds) and temprature *T* for a single RBC *N* = 1, *r* = 0.1*m* and *L* = 0.1*m*. (b) The sedimentation displacement (*x*(*m*)) of RBC as a function of time *t* (in seconds) and temprature *T* for fixed *N* = 1, *r* = 0.1*m* and *L* = 0.15*m*.

On the contrary, low-speed centrifugation leads to insufficient separation of plasma and serum from cellular blood components. By analyzing Eq. (19), one can also deduce that when the RBC form rouleaux (as *N* increases), the velocity and the position of the particle step. This is because as *N* steps up, the centrifugal force increases. This also implies since the RBC in serum forms aggregates due to clotting factors, it needs less centrifugation time than plasma. As shown in Fig. 1, the size of WBC is considerably large in comparison with the RBC. Since the platelet has the smallest size, its velocity and displacement along the test tube are significantly small. As depicted in Fig. 5a, the velocity for platelets is considerably lower than red blood cells due to the small size of platelets. Moreover, as shown in Fig. 5b, the RBC moves far more than platelets showing that to avoid the effect of clotting factors, one has to increase the centrifugation time.

**FIG. 5:**
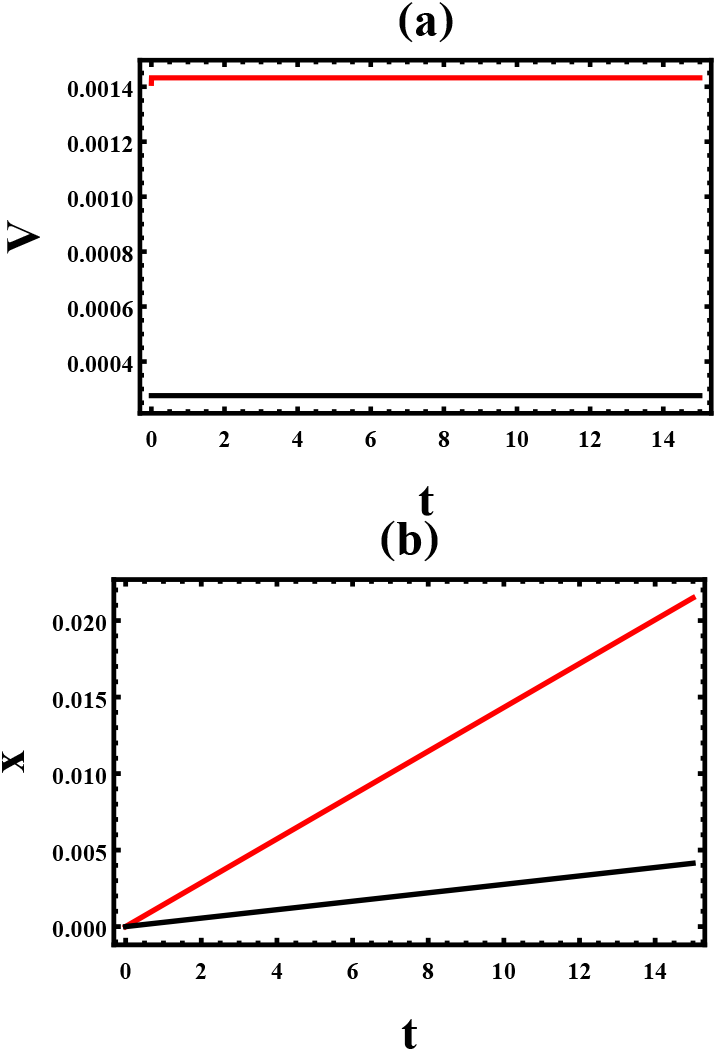
(a) The sedimentation velocity *V* of RBC (red line) and platelet (black line) as a function of time *t*. (b) The displacement of RBC (red line) and platelet (black line) as a function of time *t*. In the figures, we fix *T* = 20 degree celsius, *ω* = 200 *rad/s, r* = 0.5*m* and *L* = 0.15*m*.

## IV. THE DYNAMICS OF WHOLE BLOOD DURING CAPILLARY ACTION

In this section, we study the dynamics of the whole blood during capillary action where in this case the blood flows upward in narrow spaces without the assistance of external forces. Previous investigations show that the height that the fluid rises increases as the surface tension steps up. The viscosity of the fluid also affects the capillary action but to date, the dependence of the height on viscosity has never been explored due to the lack of mathematical correlation between the viscosity of blood and surface tension [16]. In the past, for non-Newtonian fluids, Pelofsky [17] has studied the relation between surface tension and viscosity. His empirical equation depicts that the surface tension increases as the viscosity steps up. In this work, we first analyzed the correlation between surface tension and viscous friction via data fitting.

Experimentally, the dependence of surface tension of blood on temperature was measured by Rosina *et. al*. [43]. Accordingly, the surface tension of blood is a function of temperature *T* and it is given by [16]

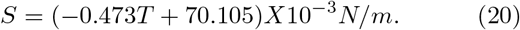

Eq. 20 exhibits that the surface tension of blood declines as the temperature increases. As discussed before, the dynamical viscous friction of blood depends on the temperature as

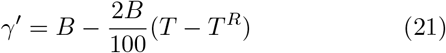

where *B* = 4*X*10^−3^*kg/ms* is the dynamical viscosity of blood at a room temperature (*T*^*R*^ = 20 degree Celsius) and *T* is the temperature [34]. The correlation between surface tension can be inferred from data fitting. Via Eqs. (20) and (21), the dependence of *S* and *γ*^*′*^ on *T* is explored as shown in Table I. Since the experiments show the surface tension *S* to have functional dependence on *γ*^*′*^, by fitting the data (see Fig. 6) depicted in Table I, one finds

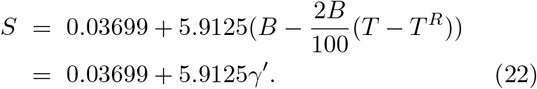

Furthermore, the surface tension is responsible for a liquid to rise up in the test tube against the downward gravitational force. As a result of the capillary action, the fluid rises to the height

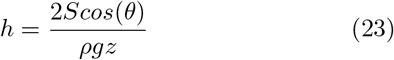

where *S* is the surface tension of the blood, *ρ* = 1060*kg/m*^3^ is the density of the fluid, *g* = 9.8*m/s*^2^ is the gravitational acceleration and *z* is the radius of the test tube. *θ* is the contact angle between the blood and the test tube while *h* is the height that the blood rises up in the test tube. For the case where *θ* = 0, we rewrite Eq. (23) as

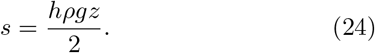

From Eqs. (22) and (24), after some algebra we get

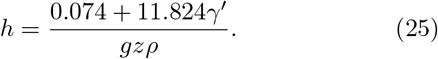

Equation (25) exhibits that the height *h* is a function of test tube radius *z* and the density of the fluid *ρ*. In a clinical laboratory, the haematocrit tube has a height of 75*mm* and inner diameter *z* = 0.4*mm*. Substituting these parameters in Eq. (25), the height that the blood rises is found to be 28*mm* at 20 degree celsius. The height *h* that the blood rises in the capillary tube is also sensitive to temperature. Via Eqs. (21) and (25), one can see that as the room temperature and inner diameter of the test tube increases, *h* decreases (see Fig. 7).

**TABLE I:**
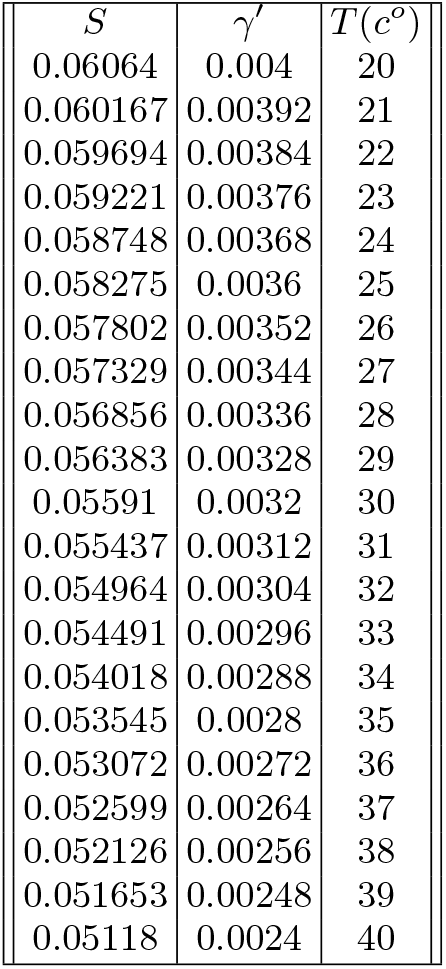
Data that show the dependence of surface tension *S* and viscous friction *γ*^*′*^ on *T*.

| $S$ | $\gamma'$ | $T(^{\circ}C)$ |
| --- | --- | --- |
| 0.06064 | 0.004 | 20 |
| 0.060167 | 0.00392 | 21 |
| 0.059694 | 0.00384 | 22 |
| 0.059221 | 0.00376 | 23 |
| 0.058748 | 0.00368 | 24 |
| 0.058275 | 0.0036 | 25 |
| 0.057802 | 0.00352 | 26 |
| 0.057329 | 0.00344 | 27 |
| 0.056856 | 0.00336 | 28 |
| 0.056383 | 0.00328 | 29 |
| 0.05591 | 0.0032 | 30 |
| 0.055437 | 0.00312 | 31 |
| 0.054964 | 0.00304 | 32 |
| 0.054491 | 0.00296 | 33 |
| 0.054018 | 0.00288 | 34 |
| 0.053545 | 0.0028 | 35 |
| 0.053072 | 0.00272 | 36 |
| 0.052599 | 0.00264 | 37 |
| 0.052126 | 0.00256 | 38 |
| 0.051653 | 0.00248 | 39 |
| 0.05118 | 0.0024 | 40 |

**FIG. 6:**
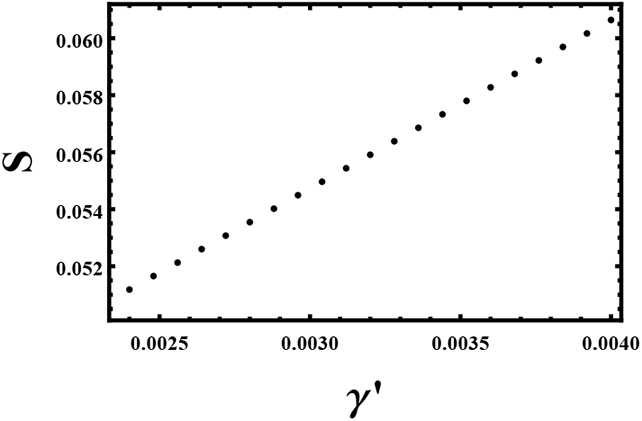
The surface tension *S* of the blood as a function of *γ*^*′*^. By fitting the data, one gets *S* = 0.03699 + 5.9125*γ*^*′*^.

**FIG. 7:**
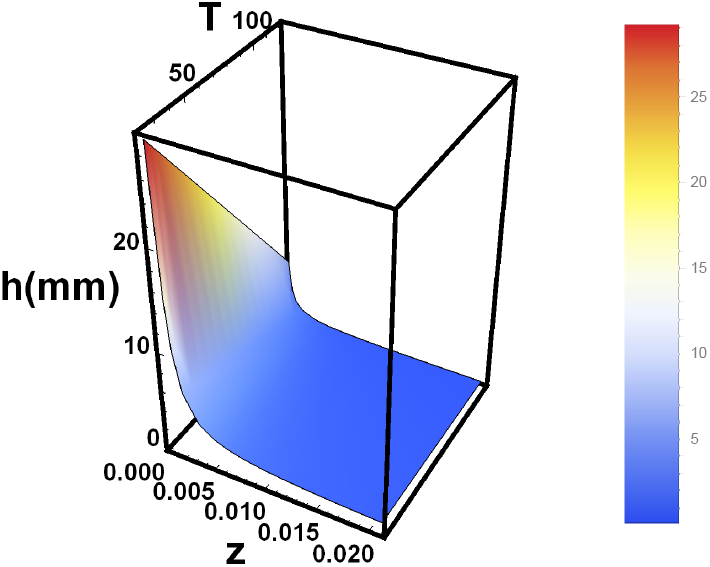
The height *h* that the blood rises in the capillary tube as a function of temperature *T* and test tube dimeter *z* for fixed-parameter *ρ* = 1060*kg/m*^3^.

The above analysis points that the increase in the room temperature results in a false positive or negative result. As depicted in Fig. 7, up to 8mm change in sedimentation can be observed when the temperature of the room varies from 20 to 40 degree celsius. When the temperature increases, the height that the blood rises decreases which is feasible since when temperature increases, the surface tension or viscosity decreases. This result also agrees experimentally. The in vivo experiments by Shinozaki *et. al*. [18] indicate that as the temperature inclines, the blood refill time decreases. The experimental observation by She *et. al*. shows that increasing temperature results in lower capillary pressures (lower capillary height) [19]. Note that the shape of RBC, plasma viscosity, and inclination of the test tube also affect the magnitude of surface tension. In anemic patients, a low hematocrit level is observed, and consequently the surface tension or viscosity of the blood decreases which in turn results in lower *h*. Excessive use of anticoagulants results in lower *h*. This is because adding too much anticoagulant decreases the viscosity or surface tension of the blood.

## V. THE ROLE OF SURFACE TENSION ON ERYTHROCYTES SEDIMENTATION RATE AND THE DYNAMICS OF BLOOD CELLS DURING CENTRIFUGATION

In this section, we explore how the surface tension of the blood affects the mobility of RBC by solving the model system analytically.

### Case 1: The effect of surface tension on the erythrocyte sedimentation rate

The erythrocyte sedimentation rate (ESR) often measures how fast a blood sample sediments along a test tube in one hour in a clinical laboratory as shown in Fig. 11. By mixing whole blood with an anticoagulant, the blood is placed in an upright Wintrobe or Westergren tube and allowed to sediments for an hour. In the Westergren method, the test tube is 200*mm* long while in the Wintrobe method the tube is only 100*mm* long. To investigate the effect of surface tension, let us rewrite Eq. (22) as

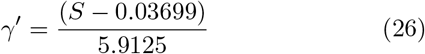

and after some algebra we get

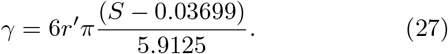

Substituting Eqs. (1) and (27) in Eq. (11), one gets

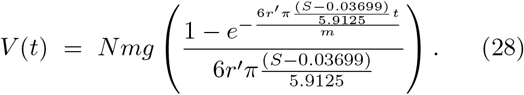

The position of the cell is then given by

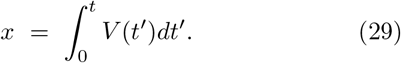

From Eqs. (28) and (29), it is evident that as the surface tension steps up, the velocity and displacement of the cells step down. Since the surface tension (Eq. 20) depends on temperature, the dynamics of the cells are also affected by the room temperature. Exploring Eqs. (28) and (29), one can see that the velocity of the cell is significantly affected by the surface tension, and time as shown in Fig. 8a. As the surface tension steps up, the velocity increases. The plot of the position of the cell as a function of time is depicted in Fig. 8b. The figure exhibits that the RBC has a faster sedimentation rate when the surface tension is smaller. From Eqs. (28) and (29), one can also see that as the RBC forms rouleaux (as *N* increases) the sedimentation rate increases. Figure 9 depicts the plot of ESR as a function of the number of RBCs and surface tension. As shown in the figure, the ESR increases as *N* increases and when the surface tension decreases.

**FIG. 8:**
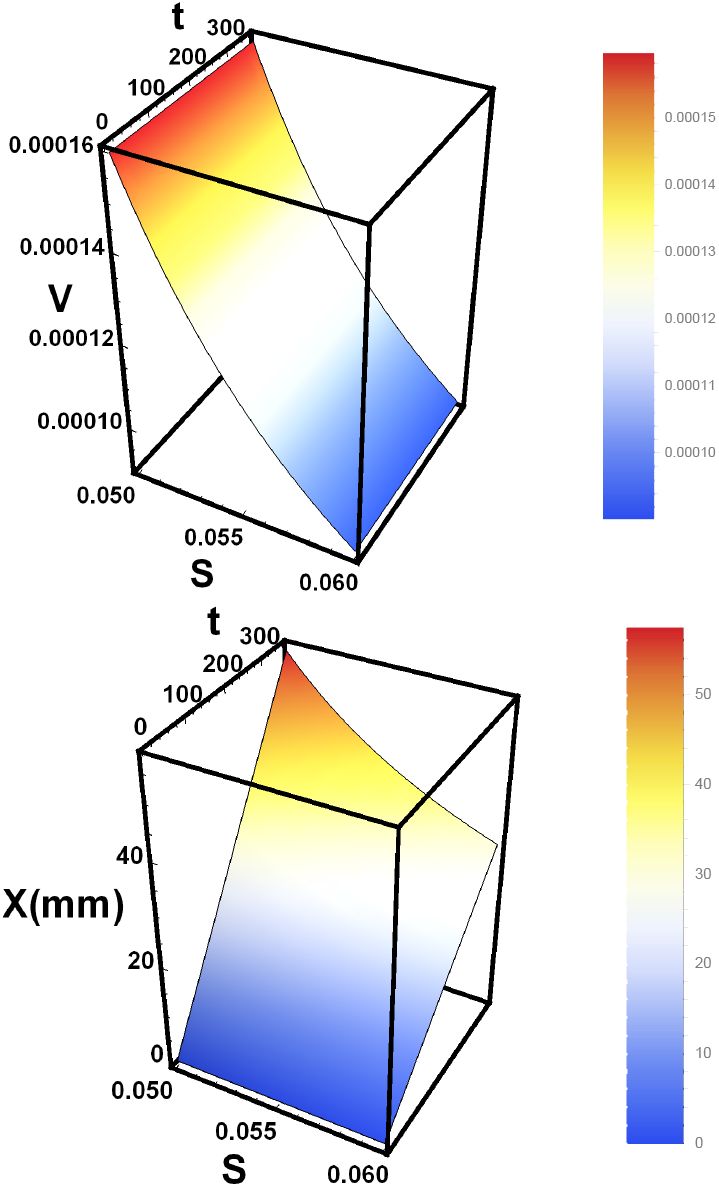
(a) The velocity (*V*) of cell as a function of *S* and *t* for *N* = 100. (b) The displacement *x* as a function of *S* and *t* for *N* = 100.

**FIG. 9:**
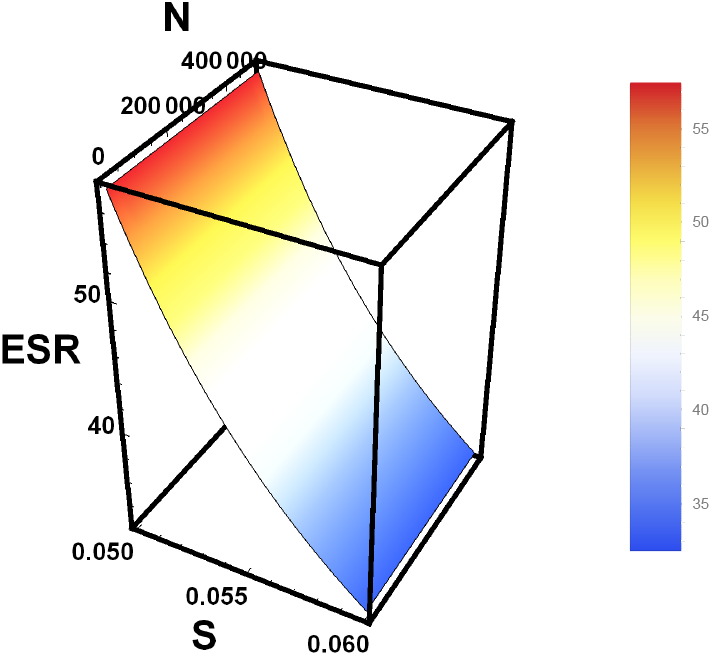
Erythrocyte sedimentation rate in one hour as a function of *N* and *S*. The ESR steps up as the number of red blood cells that form rouleaux increases and it decreases as the surface tension increases.

The background temperature of the fluid also affects the viscous friction of the fluid. As temperature increases, the viscous friction decreases, and on the contrary, the diffusibility of the cells increases as depicted in the works [30, 41]. For highly viscous fluid such as blood, when the temperature of the blood sample increases by 1 degree celsius, its viscosity steps down by 2 percent. Exploiting Eqs. (28) and (29), the dependence of the erythrocyte sedimentation rate as a function of surface tension is depicted in Fig. 10. In the figure, the number of RBCs that form clusters is fixed as *N* = 500, *N* = 600, and 800 from bottom to top, respectively. The figure depicts that the ESR decreases as the number of red blood cells (*N*) increases. When the surface tension decreases, the ESR also increases. Our analysis also shows that up to 3mm/hr sedimentation rate increase can be observed when the temperature of the room varies from 20 to 45 degree celsius. This agrees with the experimental work of Manley [44]. The ESR experiment performed via the Westergren method confirms that the increase in room temperature may significantly step up the sedimentation rate.

**FIG. 10:**
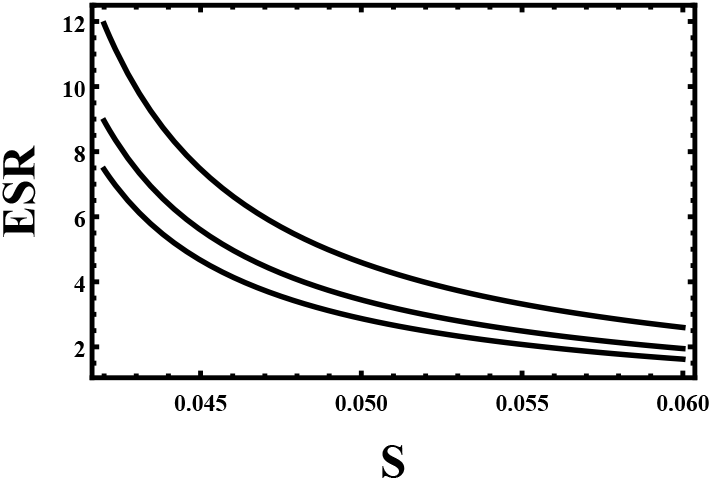
Erythrocyte sedimentation rate in one hour as a function of surface tension. The number of RBCs that form clusters is fixed as *N* = 500, *N* = 600, and *N* = 800 from top to bottom, respectively. The ESR steps up as the number of red blood cells (*N*) that form rouleaux increases as well as when the surface tension steps up.

**FIG. 11:**
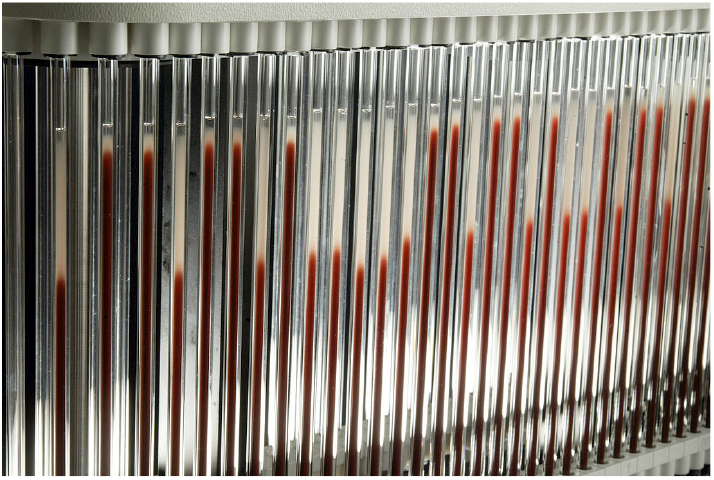
Figure that shows the sedimentation rate of RBC on Westergren pipet [42]. This hematology test is performed by mixing whole blood with an anticoagulant. The blood is then placed in an upright Wintrobe or Westergren tube. The sedimentation rate of the red blood cells is measured in millimeters (mm) at the end of one hour.

### Case 2: The role of surface tension on the dynamics of blood cells during centrifugation

As discussed before, blood cells are microscopic and they undergo a random motion in vitro when there is no external force exerted on them. Since Blood is a highly viscous medium, the chance for the RBC to accelerate is negligible even in the presence of external force. In the presence of external force (centrifugal forces), these cells undergo one-directional motion and they separate from the serum or plasma as the angular speed increases. The surface tension also affects the dynamics of the blood cells. This can be appreciated by substituting Eqs. (2) and (27) in Eq. (11). After some algebra one gets

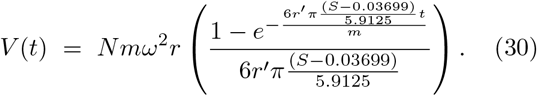

The position of the cell is then given by 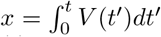. Exploiting Eq. (30), one can see that *V* (*t*) decreases as the surface tension increases (see Fig. 12). As the angular velocity increases, the velocity of the cells increases as shown in Fig. 12.

**FIG. 12:**
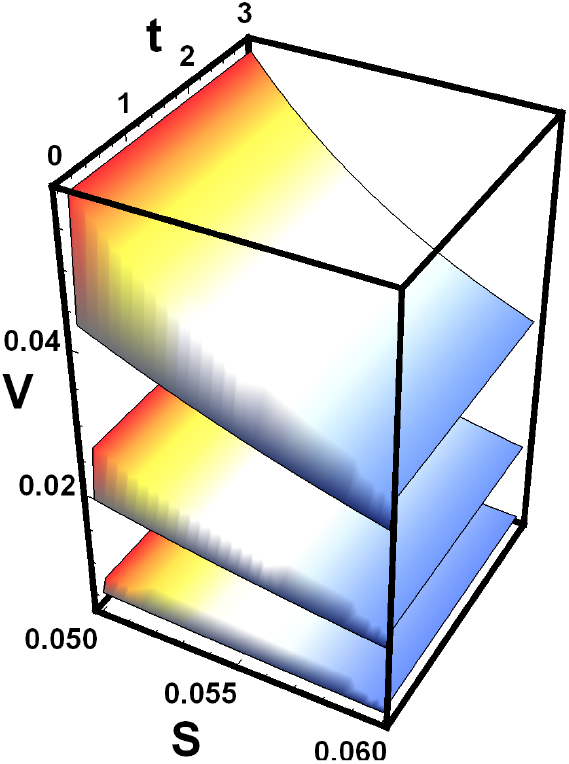
The velocity *V* as a function of time *t* and *S* for *ω* = 200*rad/s, ω* = 400*rad/s* and *ω* = 600*rad/s*, from bottom to top.

Moreover, the erythrocyte sedimentation rate depends on how the test tube is positioned. An inclination of the test tube by 3 degrees increases the ESR up to 30 percent [20]. Since blood is a highly viscous fluid, its viscosity is strongly related to the surface tension as shown in Eq. (22). On the other hand, the surface tension has a functional dependence on the degree of tilt. When the test tube becomes tilted, the surface tension of the blood decreases, and consequently its viscosity decreases. As a result a higher ESR is observed.

## VI. DISCUSSION AND RESULT

Particularly, centrifugation is one of the commonly performed laboratory procedures that help to separate blood cells such as *RBCs, WBCs*, and platelets from plasma or serum. When blood is first mixed with an anticoagulant and allowed to be centrifuged for a few minutes, the blood cells sediment by leaving the plasma at the top. In the absence of an anticoagulant, the blood clots, and when it is centrifuged, the blood cells sediment leaving the serum at the top. The serum is an ideal sample for diagnostic tests since it lacks leukocytes, erythrocytes, platelets, and other clotting factors. Although centrifugation is a routine procedure in most medical laboratories, the factors that affect the efficacy of the centrifugation process have never been studied analytically. As discussed in the work [1], the centrifugation time, temperature, the length of the test tube, and the speed of the centrifuge are the determinant factors with regard to the efficacy of the centrifugation process. In this paper, via an exact analytical solution, we study the factors that affect the efficacy of the centrifugation process. First, we examine the effect of the centrifugation time on the efficacy of the centrifugation process. Since blood cells are microscopic, their dynamics can be modeled as a Brownian particle walking in a viscous medium. As blood is a highly viscous medium, the chance for the blood cells to accelerate is negligible and the corresponding dynamics can be studied via Langevin equation or Fokker Planck equation [29–35]. Solving the Fokker-Planck equation analytically, we explore how the dynamics of blood cells behave as a function of the model parameters. Because our study is performed by considering real physiological parameters, the results obtained in this work non only agree with the experimental observations but also help to understand most hematological experiments that are conducted in vitro. In a medical laboratory, the standard test tube has a length of *L* = 150 *mm* and the cells separate from the plasma or serum at the result of the centrifugal force *f* = *Nmω*^2^*r* where *N* denotes the number of cells that form a cluster while *m* designates the mass of *RBC* or platelets. Since the *WBCs* are heavy, they move way before *RBCs*, and as an approximation, one can disregard their dynamics.

Our result depicts that the speed of the centrifuge is one of the determinant factors concerning the efficacy of the centrifugation process. As the angular speed increases, the centrifugal force steps up and as result, the particles are forced to separate from the plasma or serum. Depending on the size and fragility of the sample, the centrifugation speed should be adjected. To increase the efficacy of the centrifugation process, as the size of the particle decreases, the centrifugation speed should step up. The length of the test tube affects the effectiveness of the centrifugation process. As the length of the test, tube steps up, the centrifugal force increases. The room temperature considerably affects the dynamics of analyse during centrifugation. Most importantly, the generation of heat during centrifugation steps up the temperature within a centrifuge, and as a result, not only the stability of the sample but also the mobility of analyse is affected. The effect of temperature near the test tube causes difficulties in the current experimental procedure by inducing additional false-positive results. In this regard, developing a mathematical model and exploring the model system using the powerful tools of statistical mechanics provides insight as well as guidance regarding the dynamics of RBCs in vitro. Our result shows that as the centrifuge temperature steps up, the velocity of the cells as well as the average distance of the cell in the fluid increases. This effect of temperature can be corrected by increasing the centrifugation time. However, a longer centrifugation time might lead to a considerable amount of heat generation. This, in turn, may cause the red blood cells to lyse. Prolonged centrifugation at high speed also causes structural damage to the cells and as a result, hemolysis occurs. On the contrary, low-speed centrifugation leads to insufficient separation of plasma and serum from cellular blood components.

Our analysis also indicates that when the RBC forms rouleaux (as *N* increases), the velocity of the cells increases. This is because as *N* steps up, the centrifugal force increases. This also implies since the cells in serum form aggregates due to clotting factors, they need less centrifugation time than plasma. As shown in Fig. 1, the size of the WBC is considerably large in comparison with the RBC. Since the platelet has the smallest size, its velocity and displacement along the test tube are significantly small. As a result, the RBCs separate far more than platelets showing that to avoid the effect of clotting factors, one has to increase the centrifugation time.

Furthermore, we study the dynamics of the whole blood during capillary action where in this case the blood flows upward in narrow spaces without the assistance of external forces. Previous investigations show that the height that the fluid rises increases as the surface tension steps up. The viscosity of the fluid also affects the capillary action but to date, the dependence of the height on viscosity has never been explored due to the lack of a mathematical correlation between the viscosity of blood and surface tension [16]. In the past, for non-Newtonian fluids, Pelofsky [17] has studied the relation between surface tension and viscosity. His empirical equation depicts that the surface tension increases as the viscosity steps up. In this work, we first examine the correlation between surface tension and viscous friction via data fitting. We show that the viscosity of the blood increases as the surface tension increases. The mathematical relation between height and viscous friction is also derived. It is shown that the height that the blood rises increases as the viscous friction steps up. As the temperature of the room steps up, the height also decreases.

Moreover, in this work, we also explored the dependence of the erythrocyte sedimentation rate (ESR) on model parameters. As discussed in our recent paper [10], the erythrocyte sedimentation rate (ESR) often measures how fast a blood sample sediments along a test tube in one hour in a clinical laboratory. This analysis is performed by mixing whole blood with an anticoagulant. The blood is placed in an upright Wintrobe or Westergren tube and allowed to sediments for an hour. The normal values of the ESR vary from 0-3mm/hr for men and 0-7 mm/hr for women [21–23]. High ESR is associated with diseases that cause inflammation. In the case of inflammatory disease, the blood level of fibrinogen becomes too high [23, 24]. The presence of fibrinogen forces the RBCs to stick to each other and as a result, they form aggregates of RBC called rouleaux. As the mass of the rouleaux increases, the weight of the rouleaux dominates the vicious friction and as a result, the RBCs start to precipitate. The temperature of the laboratory (blood sample) also significantly affects the test result [25]. As the temperature of the sample steps up, the ESR increases. In the past, a mathematical model was developed by Sharma .*et. al* to study the effect of blood concentration on the erythrocyte sedimentation rate [26]. Later the effect of the concentration of nutrients on the red blood cell sedimentation rate was investigated in the work [27]. More recently, the sedimentation rate of RBC was explored via a model that uses Caputo fraction derivative [28]. The theoretical work obtained in this work was compared with the sedimentation rate that was analyzed experimentally. All of these experimental and theoretical works exposed the factors that affect the sedimentation rate of RBCs. In this paper, extending our recent work [10], we study how surface tension as well as the tilt in test tube angle affects the ESR.

## VII. SUMMARY AND CONCLUSION

In this paper, via an exact analytical solution, we study the factors that affect the efficacy of the centrifugation process. The effect of the centrifugation time on the efficacy of the centrifugation process is explored by studying the dynamics of the blood cells via the well-known Langevin equations or equivalently, by solving Fokker-Plank equations. As blood cells are microscopic, their dynamics can be modeled as a Brownian particle walking in a viscous medium. Since blood is a highly viscous medium, the chance for the blood cells to accelerate is negligible and the corresponding dynamics can be studied via the Langevin equation or Fokker-Planck equation. Solving the Fokker-Planck equation analytically, we explore how the dynamics of blood cells behave as a function of the model parameters. Because our study is performed by considering real physiological parameters, the results obtained in this work not only agree with the experimental observations but also help to understand most hematological experiments that are conducted in vitro or in vivo.

In a medical laboratory, the standard test tube has a length of *L* = 150*mm* and the cells separate from the plasma or serum at the result of the centrifugal force *f* = *Nmω*^2^*r* where *N* denotes the number of cells that form a cluster while *m* designates the mass of *RBC* or platelets. Since the *WBCs* are heavy, they move way before *RBCs*, and as an approximation, we disregard their dynamics.

The speed of the centrifuge is one of the main factors concerning the efficacy of the centrifugation process. It is shown that as the angular speed increases, the centrifugal force steps up and as a result, the particles are forced to separate from the plasma or serum. Based on the size and fragility of the sample, the centrifugation speed should be adjected. To increase the efficiency of the centrifugation process, as the size of the particle decreases, the centrifugation speed should step up. For instance, our work depicts that the velocity for platelets is considerably lower than red blood cells due to the small size of platelets. The RBC separates far more than platelets showing that to avoid the effect of clotting factors, one has to increase the centrifugation time. We also show that as the length of the test tube steps, the centrifugal force increases. The dynamics of the cells during centrifugation is also affected by the room temperature. The generation of heat during centrifugation steps up the temperature within a centrifuge and as a result, not only the stability of the sample but also the mobility of the sample is affected. Our result shows that as the centrifuge temperature steps up, the velocity of the cells as well as the average distance of the cell in the fluid decreases. This effect of temperature can be corrected by increasing the centrifugation time. However, a longer centrifugation time might lead to a considerable amount of heat generation. This, in turn, may cause the red blood cells to lyse. Prolonged centrifugation at high speed also causes structural damage to the cells and as a result, hemolysis occurs. On the contrary, low-speed centrifugation leads to insufficient separation of plasma and serum from cellular blood components. When the RBC forms rouleaux (as *N* increases), the velocity and the position of the particle step up. This is because as *N* increases, the centrifugal force increases. This also implies since the RBC in serum forms aggregates due to clotting factors, it needs less centrifugation time than plasma.

The dynamics of the whole blood during capillary action is studied where in this case the blood flows upward in narrow spaces without the assistance of external forces. In this work, we first analyzed the correlation between surface tension and viscous friction via data fitting. We show that the viscosity steps up linearly as the surface tension increases. The mathematical relation between height and viscous friction is derived. It is shown that the height of the blood rises increases as the viscous friction increases. As the temperature of the room steps up, the height decreases. The dependence of the erythrocyte sedimentation rate (ESR) on model parameters is also studied.

Most medical laboratory examinations are affected by external factors such as temperature. To understand the factors that affect the outcome of these routine examinations, it is vital to explore the dynamics of the whole blood, erythrocytes (RBCs), leukocytes (WBCs), and thrombocytes (platelets). Since our mathematical analysis is performed by using physiological parameters, all of the results depicted in this work can be reconfirmed experimentally. The simplified model presented in this work can also help to understand most hematological experiments that are conducted in vitro or in vivo.

## Acknowledgment

I would like to thank Mulu Zebene and Blyanesh Bezabih for the constant encouragement.

## Author contribution statements

Mesfin Taye conceived the research idea, developed the theory, and performed the analytical computations. He also contributes to the writing of the manuscript.

